# Chitosan stimulates root hair callose deposition and inhibits root hair growth

**DOI:** 10.1101/2023.07.30.551171

**Authors:** Matěj Drs, Samuel Haluška, Eliška Škrabálková, Pavel Krupař, Andrea Potocká, Lucie Brejšková, Karel Muller, Natalia Serrano, Aline Voxeur, Samantha Vernhettes, Jitka Ortmannová, George Caldarescu, Matyáš Fendrych, Martin Potocký, Viktor Žárský, Tamara Pečenková

**Affiliations:** Institute of Experimental Botany of the Czech Academy of Sciences, Rozvojová 263, 165 02, Prague 6, Czech Republic; Department of Experimental Plant Biology, Faculty of Science, Charles University, Viničná 5, 128 44, Prague 2, Czech Republic; Université Paris-Saclay, INRAE, AgroParisTech, Institut Jean-Pierre Bourgin (IJPB), 78000, Versailles, France

## Abstract

Although angiosperm plants have a general capacity to react after the immunity elicitor chitin or chitosan treatment by the cell wall callose deposition, this response in particular cell types and its evolutionary conservation is not understood. Here we show that also the growing root hairs (RHs) of Arabidopsis can respond to a mild (0.001%) chitosan treatment by the callose deposition and by a deceleration of the RH growth. We demonstrate that the glucan synthase-like 5 (GSL5)/PMR4 is vital for chitosan-induced callose deposition but not for RH growth inhibition. Upon the higher chitosan concentration (0.01%) treatment, RHs do not deposit callose, while growth inhibition is prominent. To understand the specificities of the low and high concentration chitosan treatments, we analysed the corresponding PTI signalling components, gene expression, and RH cellular endomembrane and cytoskeleton modifications. Importantly, chitosan-induced callose deposition is also present in the functionally analogous and evolutionarily only distantly related RH-like structures rhizophores (lycophytes) and rhizoids (bryophytes). Our results point to the RH callose deposition as a conserved strategy of soil-anchoring plant cells (rhizoids/rhizophores/RHs) to deal with mild biotic stress. At the same time, high chitosan concentration prominently disturbs intracellular dynamics, tip-localised endomembrane compartments and RH growth, precluding callose deposition.

## Introduction

Root hairs (RHs) are unicellular plant structures crucial for several aspects of root function: mechanical anchorage of a plant in a substrate, the absorption of water and essential nutrients due to increased root surface, and in root-microbe interactions. In the model plant *Arabidopsis thaliana*, RH cell fate is determined by a position-dependent signalling cascade governed by specific transcription factors, leading to the RH initiation, bulging and cylindrical/polarised tip growth (Carol and Dolan, 2002; Dolan, 1996; Parker et al., 2000; Vissenberg et al., 2020). Several root hair-defective (RHD) genes, F-actin cytoskeletal meshwork and secretory pathway play a prominent role in this process (Baluška et al., 2000; Ichikawa et al., 2014). The tip growth is supported by constant modification of the preexisting cell wall (CW), as documented by the vital importance of several cellulose and hemicellulose biosynthesis participants (Cavalier et al., 2008; Park et al., 2011).

In aerial plant parts, when a pathogen attacks the plant cell, pathogen-associated molecular patterns (PAMP) triggered immunity (PTI) and potentially effector-triggered immunity (ETI) are activated (Jones and Dangl, 2006; Ngou et al., 2021). One of the earliest defence responses to a pathogen attack is the deposition of β-1,3 glucan polysaccharide callose by which the plant reinforces cell walls and blocks the pathogen entry (Jacobs et al., 2003). In Arabidopsis, among 12 callose synthase genes, only one of them, *GSL5* or *PMR4* (*POWDERY MILDEW RESISTANT 4*), is expressed upon pathogenic infection or elicitor treatment, such as fungal cell wall derived chitin and chitosan, but also endogenous damage-associated patterns (DAMP) (Denoux et al., 2008; Gómez-Gómez et al., 1999; Luna et al., 2011; Verma and Hong, 2001; Vogel and Somerville, 2000).

In contrast to shoots, the roots are constantly exposed to a mixture of microorganisms, and plants need to prevent the over-activation of immunity in roots, mainly by restricting defence responses to specific root zones and cell layers (Chuberre et al., 2018). The PTI in roots also comprises the ROS (reactive oxygen species) production, sporadic and patchy callose deposition and growth rate modifications upon PAMP stimulation (Badri et al., 2008; Chuberre et al., 2018; Millet et al., 2010; Rich-Griffin et al., 2020; Zhang et al., 2022). Upon injury of RH, callose can be deposited locally in the RH cell wall as well, however, the identity of the callose synthase involved remains unknown (Galway et al., 2011). It has been recently shown that the defence-related callose synthase PMR4 is responsible for the RH callose deposition upon phosphate starvation (Okada et al., 2023). The RH can react to pathogenic bacteria (e. g. *Pseudomonas syringae*), similarly to the beneficial ones, by the enhancement of the RH growth (Pečenková et al., 2017; Zamioudis et al., 2013). Unlike living bacteria, elicitor treatments in Arabidopsis do not provoke prominent RH growth changes, except for the DAMP PEP1 (Okada et al., 2021).

Both RH tip growth and pathogen sensing are under the control of development- and defence-related receptor kinases, and binding of their cognate ligands can for instance lead to the ROS production and/or activation of Ca2+ channels, such as CYCLIC NUCLEOTIDE GATED CHANNELS (CNGCs) (Ladwig et al., 2015)). The CNGCs produce transients with specific patterns, further transmitted to transcriptional reprogramming of defence (or symbiosis) (Yuan et al., 2017). Recent studies have reported that CNGC6, CNGC9 and CNGC14 are crucial for RH tip polar growth (Brost et al., 2019; Ladwig et al., 2015). A phosphorylation activation of mitogen-activated protein kinases (MAPK) cascade is another convergence point of signalling pathways. The MAPK3/6 signalling cascade plays a role in the RH growth (Rentel et al., 2004), as well as in plant immunity (including chitin-activated defence), together with the second MAPK4-comprising cascade (rev. in (Zhang and Zhang, 2022)).

The simplest way to study defence responses is by applying a single PAMP or immunity elicitor such as chitin or its derivate chitosan. Chitin is an insoluble polymer of β-1,4-linked N-acetylglucosamine, which is in Arabidopsis recognized by a complex of receptor and co-receptors - Chitin Elicitor Receptor Kinase 1 (CERK1) and lysin motif receptor kinases 4 and 5 (LYK4 and 5) (Cao et al., 2014; Miya et al., 2007; Wan et al., 2012). The chitin recognition activates the production of chitinases by plants; to avoid this recognition and hydrolysis by chitinases, fungi secrete chitin de-N-acetylases to convert chitin to chitosan. Fully deacetylated polymer units can still activate Ca^2+^ signalling-related subset of cellular responses, which in case of overactivation provokes cell death independent of the CERK1/ROS/MAPK pathway (Ye et al., 2020).

Chitosan is typically obtained from marine crustaceans’ chitin by chemical deacetylation (up to 90%) with molecular weights (Mw) ranging from oligo-(<5 kDa) to polysaccharides (70–100 kDa). Due to the protonation of glucosamine, chitosan molecules carry an overall positive charge. Chitosan functions as a PAMP (Iriti et al., 2006), but also as an antimicrobial factor causing intracellular ROS burst with subsequent oxidation of pathogen cell membrane fatty acids and plasma membrane permeabilization (Lopez-Moya et al., 2017). Interestingly, high doses of chitosan in the rhizosphere of Arabidopsis, tomato or barley significantly arrest root development, probably via interference with auxin and gibberellic acid regulation or via activation of jasmonic (JA) and salicylic acid (SA) related genes in roots (Lopez-Moya et al., 2017; Suwanchaikasem et al., 2022).

In this report, we compare the elicitation of plant immunity with two different chitosan concentrations, low 0.001%, further referred to as LCC, and high, 0.01% further referred to as HCC, focusing on RH reactions. We bring evidence of the plant capability to adjust the RH rate growth, as well as to reinforce the RH cell wall by the callose deposition. The observed RH-specific effects are dependent on the chitosan concentration, but not its molecular size, and are present also in the functionally analogous but evolutionary non-related structures, namely rhizoids of the liverwort *Marchantia polymorpha* and rhizophores of the lycophyte *Selaginella uncinata*, as well as in the regenerating protoplasts of *Physcomitrium patens*. Further, on the subcellular level, we tested the responsiveness of RH endomembranes and cytoskeleton to chitosan treatments. To understand the impact of the chitosan treatments on overall plant growth and defence, we performed Ca^2+^ and MAPK signalling assays and RNAseq analysis, pointing to limited but significant changes caused by LCC, compared to HCC treatment. Out of these experiments, we could conclude that the RH callose deposition is the consequence of the mild biotic stress affecting the cells with the intensive growth rate, while the stronger biotic stress provokes almost immediate and more prominent tip-localised endomembrane changes and RH growth cessation. Our results thus reveal the unexpected aspects of the RH growth-defence trade-off, and set the basis for possible future modifications of the plant fitness using mild and environmentally low-impact treatments.

## Results

### Chitosan treatment induces concentration-dependent root growth changes and callose deposition also in root hairs

When the 5-7 day-old *A. thaliana* seedlings in liquid medium were treated with several different concentrations of chitosan 0.0001% - 0.01%, 24 h post treatment we could observe the ectopic callose deposition in cotyledon leaves, but also in roots and RHs (Figure 1A). While the LCC (0.001%) induced small callose deposits in leaves, no primary root growth inhibition and prominent callose depositions in RH regions close to the root tips, the HCC (0.01%) induced larger callose deposits in leaves, prominent primary root growth inhibition and no callose in root hairs. Conversely, plants incubated with very low chitosan concentration (0.0001%) were indistinguishable from the mock treatment by both the lack of prominent callose deposition and no inhibition of the primary root growth (Figure 1A, B). Given the striking response of RH to specific chitosan concentration (LCC, 0.001%), we wondered if it is caused by the particular chitosan composition. We therefore tested the effect of the same low concentration for several other chitosan variants with different Mw, viscosity, and solubility. The observed effect on the callose deposition along RH was consistent for all analysed chitosan variants but not for the non-deacetylated chitin, confirming that the RH callose deposition is chitosan-specific (Figure 2). When the seedlings were treated by LCC in distilled water, which does not support the RH growth such as liquid ½ MS, there was no callose deposition in existing RHs, pointing to the fact that the callose is deposited only in newly growing RH exposed to chitosan (Supplementary figure 1A). Similarly, the application of HCC provokes strong RH growth inhibition without observable RH callose deposition (Figure 1, Supplementary figure 1B).

**Figure 1.**
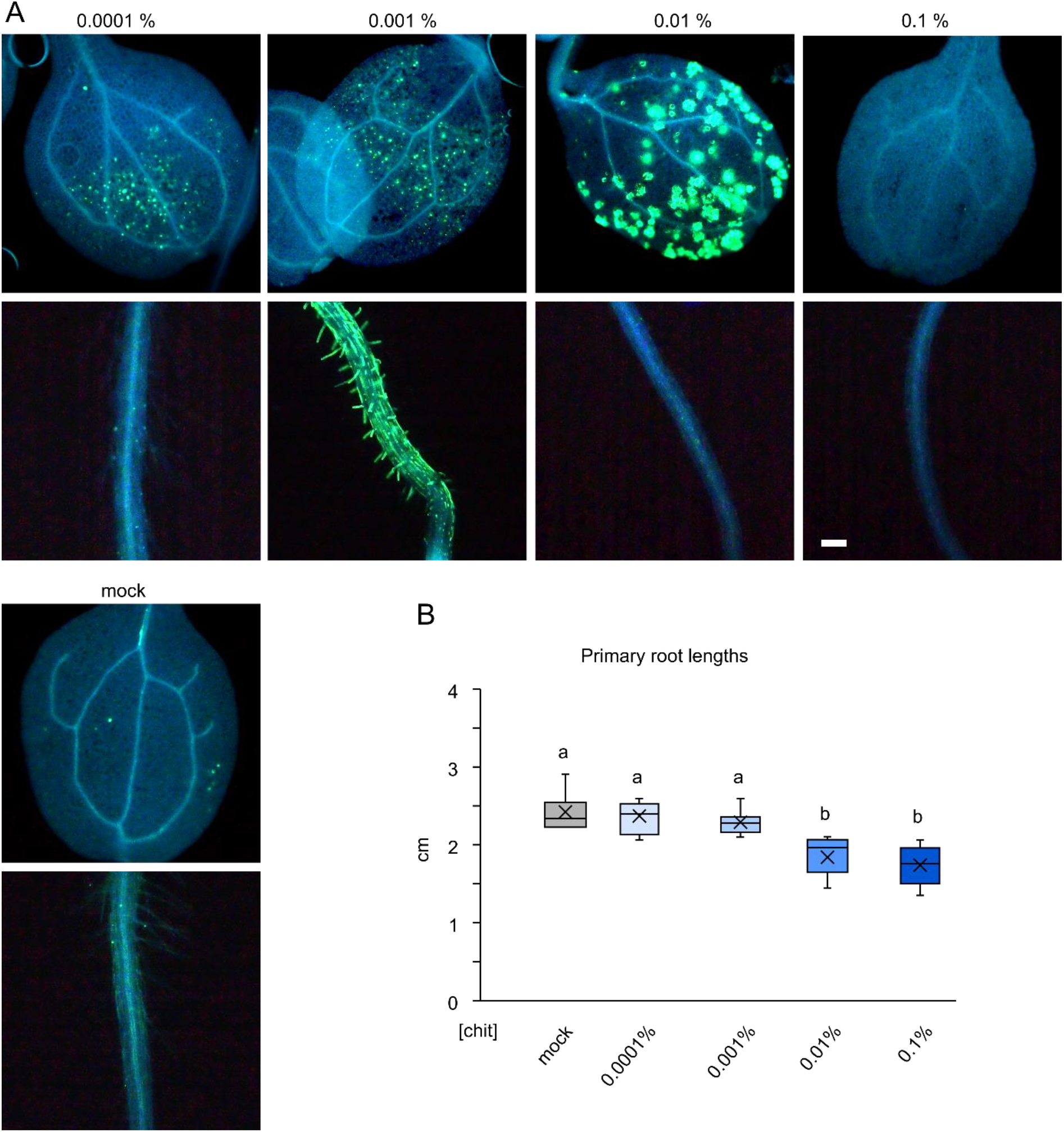
Concentration dependent modes of chitosan-triggered callose deposition. Different concentrations of chitosan cause different patterns of callose depositions 24 h post treatment (p. t.) in both cotyledons and roots (root hairs); mock treatment as a control below (bar = 100 µm). B) Concentration dependent primary root growth inhibition 24 h p. t. n=8 - 11; small letters indicate significance of differences based on the ANOVA test.

**Figure 2.**
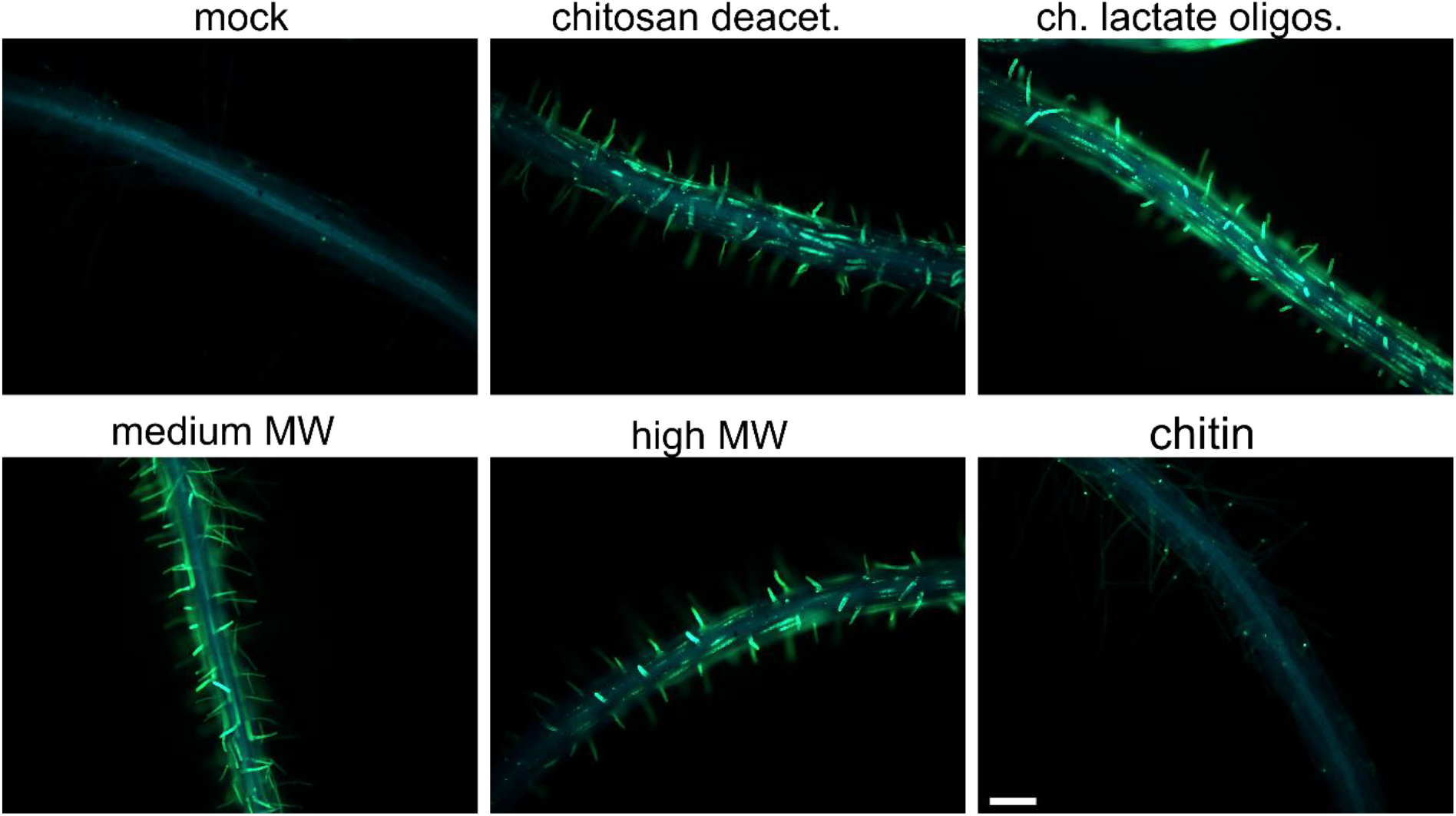
Root hair growth during the chitosan treatments. A) The low chitosan concentration (0.001%, LCC) of several other chitosan variants has the same effect on RH callose deposition (bar = 100 µm).

We could conclude that chitosan provokes specific responses in both above-ground organs and root hairs, depending on the applied concentrations.

### Callose deposition is PMR4-dependent and uncoupled from the RH growth inhibition

In order to uncover the key players in the RH callose deposition, we decided to analyse the RH modifications in the *gsl5*/*pmr4* mutant lacking the callose synthase isoform responsible for the pathogen-induced callose deposition. The LCC-induced callose deposition was completely abolished in the *pmr4* mutant (Figure 3A). Further on, when using the GFP-tagged PMR4 construct (Sabol et al., in preparation), we could observe a higher accumulation of PMR4 on the tip of the newly growing RH 4h after the chitosan addition in comparison to the mock treatment (Figure 3B). Along with the callose *pmr4* mutant, we tested *cals8-1,* which is involved in the biotic stress-related regulation of plasmodesmatal permeability, as well as a range of secretory, defence-related (including signalling co-receptor *bak1* and chitin receptor *cerk1*) and phytohormone mutants, however, for all of them we found the RH callose pattern similar to the one in WT/Col-0 (Supplementary figure 1C).

**Figure 3.**
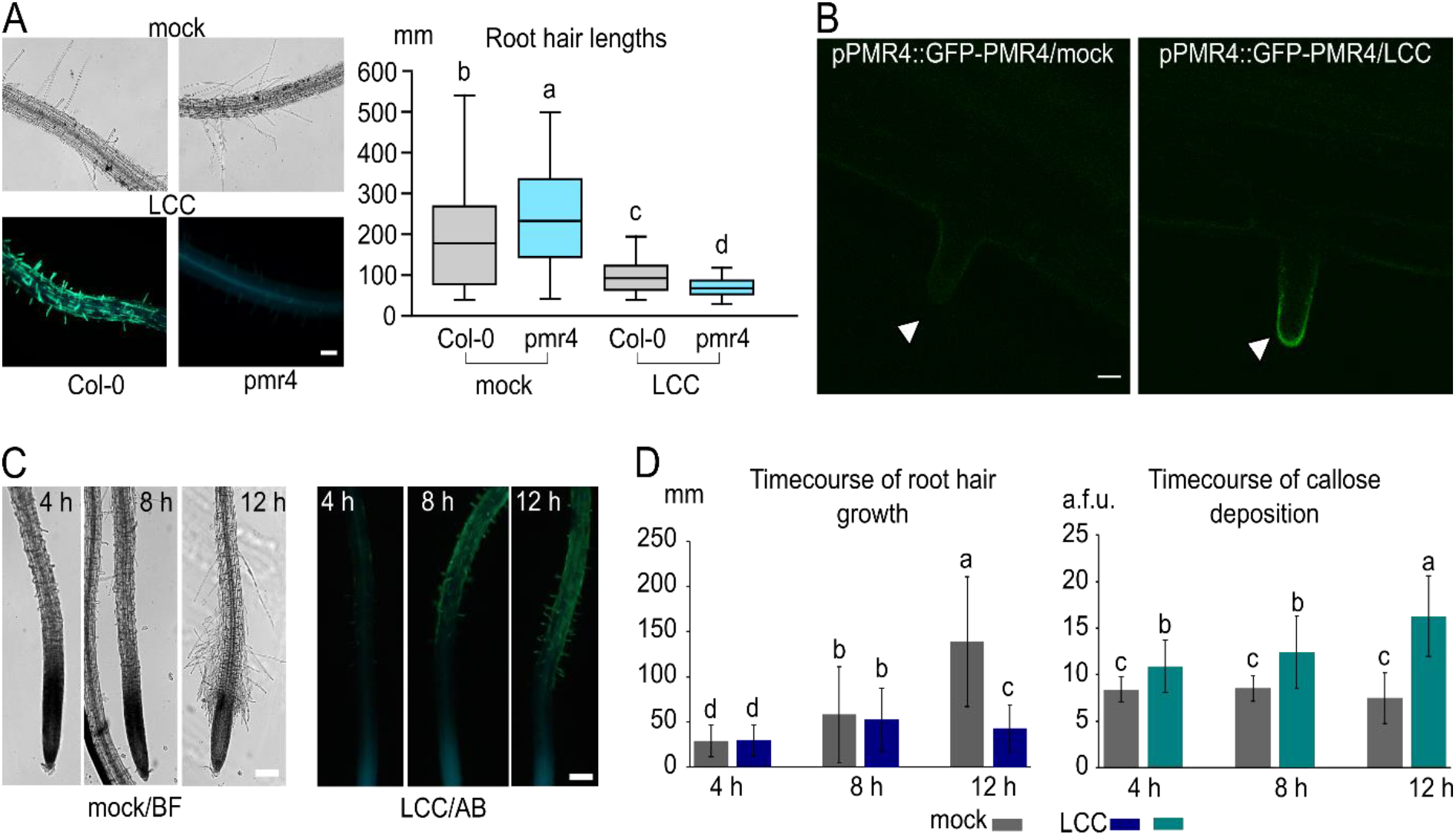
Callose deposition in LCC is PMR4-dependent. A) A *pmr4* mutant root hair (RH) growth does not deposit callose upon the LCC treatment; nevertheless, the RH growth is suppressed to the similar extent as in the Col-0 control. Small letters indicate significance of differences based on the ANOVA test; n=100-140. B) Upon the LCC treatment, at 4 h p. t. GFP-PMR4 accumulates in the tips of growing RH, further confirming a role for PMR4 in RHs callose deposition (marked by white asterisks). C) The appearance of root hairs of mock-treated (bright field, BF) and LCC-treated (callose stained with aniline blue, AB) seedlings at 4, 8 and 12 h p. t. D) Timecourse of the decline of the RH growth and the callose deposition reveals significant changes between the mock- and LCC- treated seedlings in the growth observable at 12 h p.t. (n=90-140), and in the callose deposition already at 4 h p. t. (n=20-35). Small letters indicate significance of differences based on the ANOVA test. Bars = 100 µm in A), C) and D) and 10 µm in B).

Since the LCC caused a decrease in RH growth, we hypothesised that it may be coupled with callose deposition and rigidification of the RH cell wall. However, the LCC-induced RH growth decline in the *pmr4* mutant that does not deposit callose was similar to that of WT/Col-0 (Figure 3A). We also monitored both callose deposition and root hair growth in WT after the LCC addition, showing that the callose starts to accumulate already at 4h time point, while the average inhibition of the root hair growth starts to be evident after 8h, reaching significant difference from mock treatment approximately 12 h after addition of chitosan (Figure 3C and 3D). This difference in the time course pattern further confirms the independence of the RH growth decline and callose deposition. In contrast to LCC, the HCC treatment completely arrested the growth of new RHs already at the 4h time point (Supplementary figure 1B).

The LCC-induced callose deposition in RHs is thus PMR4-dependent and uncoupled from the mechanisms underlying slower RH growth.

### Low concentration chitosan treatment triggers weak PTI-related signalling

We further wanted to uncover differences in general activation of PTI responses between the LCC and HCC treatments. For that purpose, we first followed calcium spiking using sensor R-GECO1 construct (Keinath et al., 2015) in a 20 min treatment of seedlings by the two concentrations of chitosan. There was only a mild Ca^2+^ increase in spiking in LCC treatment compared to HCC treatment, which triggered more prominent peaks of intracellular Ca^2+^ increase (Figure 4A). Besides their intensity, the two treatments also differ in their patterns and peaking time, evidencing early specificity of the LCC versus HCC. Interestingly, despite the obvious employment of the Ca^2+^ channels in chitosan sensing, *cngc* mutants have unaffected deposition of the callose to the RH cell walls, indicative of higher importance of the CNGC-mediated signalling for the RH growth than for the callose deposition (Supplementary figure 1C).

**Figure 4.**
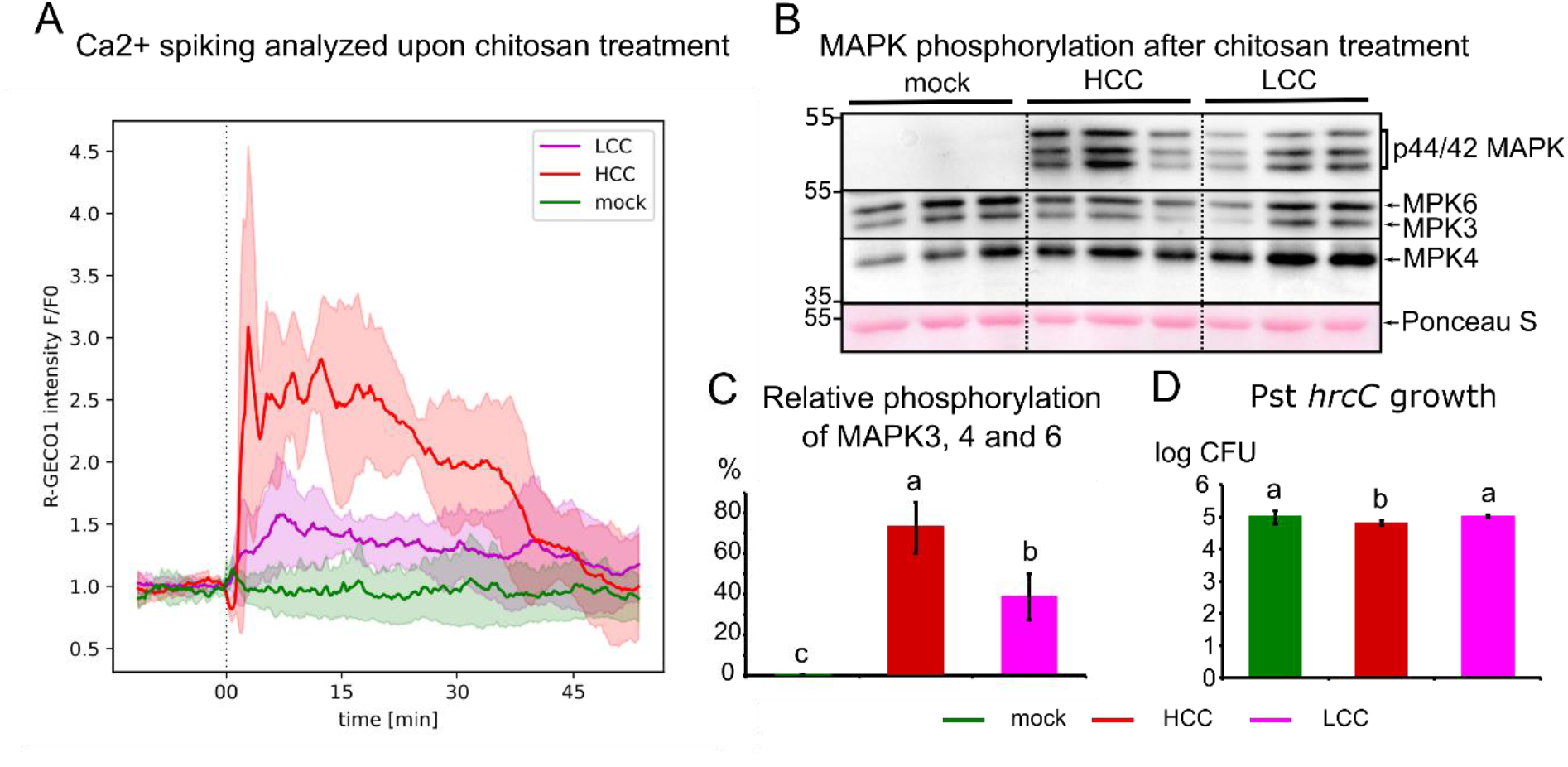
Chitosan-triggered signalling in *A. thaliana* seedlings. A) Ca^2+^ spiking analysed as R-GECO1 sensor fluorescence in root cells upon chitosan treatment shows slightly later onset with mild oscillations and long duration within the ca 45 min frame of LCC treatment, while HCC induces prominent Ca^2+^ spiking with subsequent drop in signal intensity. The error envelope represents SD, n=4-8. B) Western blot analysis of MAPK activation by phosphorylation upon 30 min of HCC and LCC treatments quantified as the ratio of band intensities obtained for phosphorylated portion versus the whole pool MAPKs. Both treatments cause significant increase in MPK3/6/4 phosphorylation in comparison to the mock treatment, also the difference between the LCC- and HCC-caused phosphorylations is significant, with LCC causing almost two-time less MAPKs phosphorylation. Small letters indicate significance of differences based on the ANOVA test, n=9. D) Bacteria Pst *hrcC* amplification in seedlings pretreated with mock, HCC and LCC (24 h of pretreatment + 24 h of bacteria inoculation); n=7-8, small letters indicate significance of differences based on the ANOVA test.

Additionally, we assayed another parallel early signalling PTI pathway, MAPK activation, after the two chitosan concentrations 30 min long treatments. Similarly, a western blot analysis of MAPK activation by phosphorylation (quantified as a ratio of band intensities obtained for the phosphorylated part and whole pool MAPKs) demonstrates significant MAPK activation in both cases. However, HCC causes a more prominent effect on the MAPK signalling pathway than the LCC (Figure 4B and C). The distribution of activation along the three tested MAPKs - MAPK3,6 and 4 was similar for both treatments.

Accordingly with these results, the pretreatment of seedlings with HCC, and not LCC, causes significant enhancement of PTI, as demonstrated by the bacteria sensitivity and amplification assay upon seedlings inoculation with *Pseudomonas syringae pv. tomato* mutant in T3 secretion system *hrcC* (Pst *hrcC*; Figure 4D).

These results confirm that the LCC treatment, similarly to the HCC one, activates PTI-related signalisation, nevertheless, the pattern and intensities of these activations significantly differ.

### Chitosan modulates rapid calcium and endomembrane dynamics but not actin cytoskeleton in growing root hairs

Having shown distinct fast calcium responses after LCC and HCC treatment, we next analysed the spatiotemporal details of the Ca^2+^ reporter R-GECO1 within the RH cells. In mock treatment, the very tips of RH undergo constitutive pulsatile changes of R-GECO1 signal intensity, while the atrichoblastś R-GECO1 pattern transiently changes upon the buffer exchange during the mock treatment, and much more prominently upon chitosan treatments (Figure 5A and B, Supplementary video). Each of the applied treatments causes specific RH Ca^2+^ influx intensities and patterns, with LCC treatment causing even a decrease in the RH-tip located cytoplasmic Ca^2+^ during the initial phase of the treatment (Figure 5C). Interestingly, upon HCC, a wave of R-GECO1 signal often starting within 1-2 min at the RH tips and moving to the RH cell interior and further on through trichoblast toward other root cells was observed (Supplementary video).

**Figure 5.**
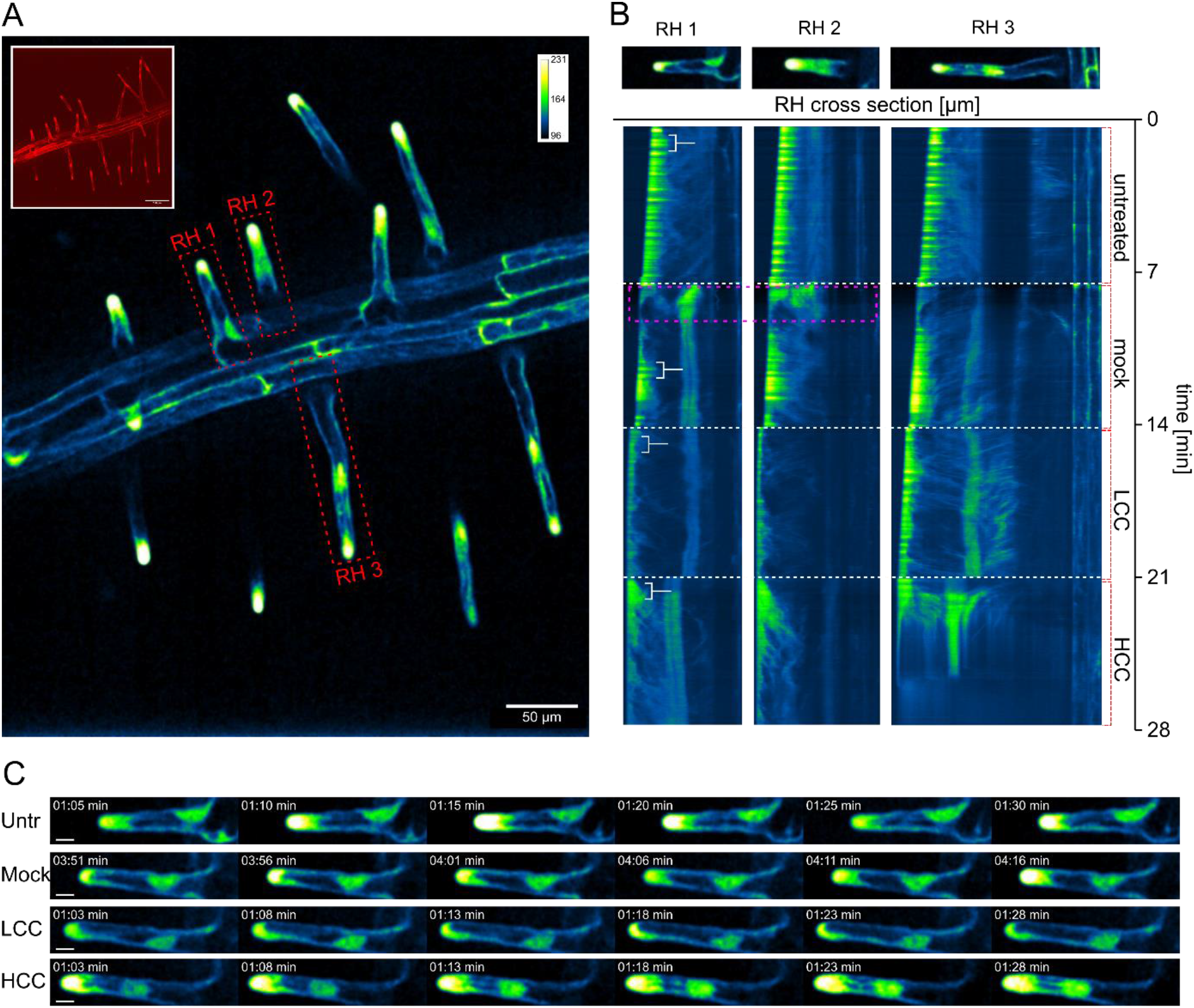
R-GECO1 fluorescence patterns in RHs of treated seedlings. (A) represents the root of 6-day-old seedling expressing R-GECO1 (an example of the red signal appearance shown in upper left box), with indicated root hairs analysed in section (B) (red dashed rectangle; RH1, RH2, RH3). Artificial LUTs (green fire blue) were used; the calibration bar represents the signal intensity (box on the right). (B) shows R-GECO1 oscillations depicted via kymograph throughout the experiment (Untreated, mock, LCC, HCC) of 3 selected RH (RH1-3). White dashed lines indicate the end of a particular part of the treatment, magenta dashed rectangle indicates the specific oscillation pattern caused by flooding the sample with ½ MS solution. White brackets indicate specific time windows used for the section (C). Section (C) represents 4 typical R-GECO1 oscillation patterns for each treatment (Untr (untreated), mock, LCC, HCC) visualised on 6 consecutive frames with 5 s gaps of RH1. Scale bar = 10 µm.

We then tested if and how the chitosan-triggered early signalling event affects the intracellular dynamics. We focused on well-growing RH close to the root tip, and after applying mock, LCC and HCC treatments, we followed intracellular changes in localisation and dynamics of vacuolar and cytoskeletal genetic markers, as well as endocytic dye FM4-64 in a real-time approach (Figure 6). We monitored treated RH for 0-30 minutes. First, we could observe an immediate and significant root hair growth cessation in the case of HCC-treated seedlings, often accompanied by RH damaging and bursting. In addition, for LCC we could also observe significantly lower average growth rates. Nevertheless, out of the obtained data distribution, we could conclude that in the case of this treatment, RHs mostly either prominently slow down their growth, or they grow at rates similar to mock-treated root hairs (Figure 6A). Interestingly, the vacuolar dynamics does not follow the trend found for growth and the RHs react to the two treatments contradictory – while LCC causes small and insignificant enhancement in vacuolar dynamics, HCC impairs the vacuolar dynamics in the RH tip area (Figure 6B). The cessation of vacuolar movement in the RH tip upon HCC treatment may be related to observation obtained for the same treatment on the enhanced FM4-64 dye accumulation in RH endocytotic/vesicular compartment, probably as a consequence of collapsing endomembrane trafficking (Figure 6C).

**Figure 6.**
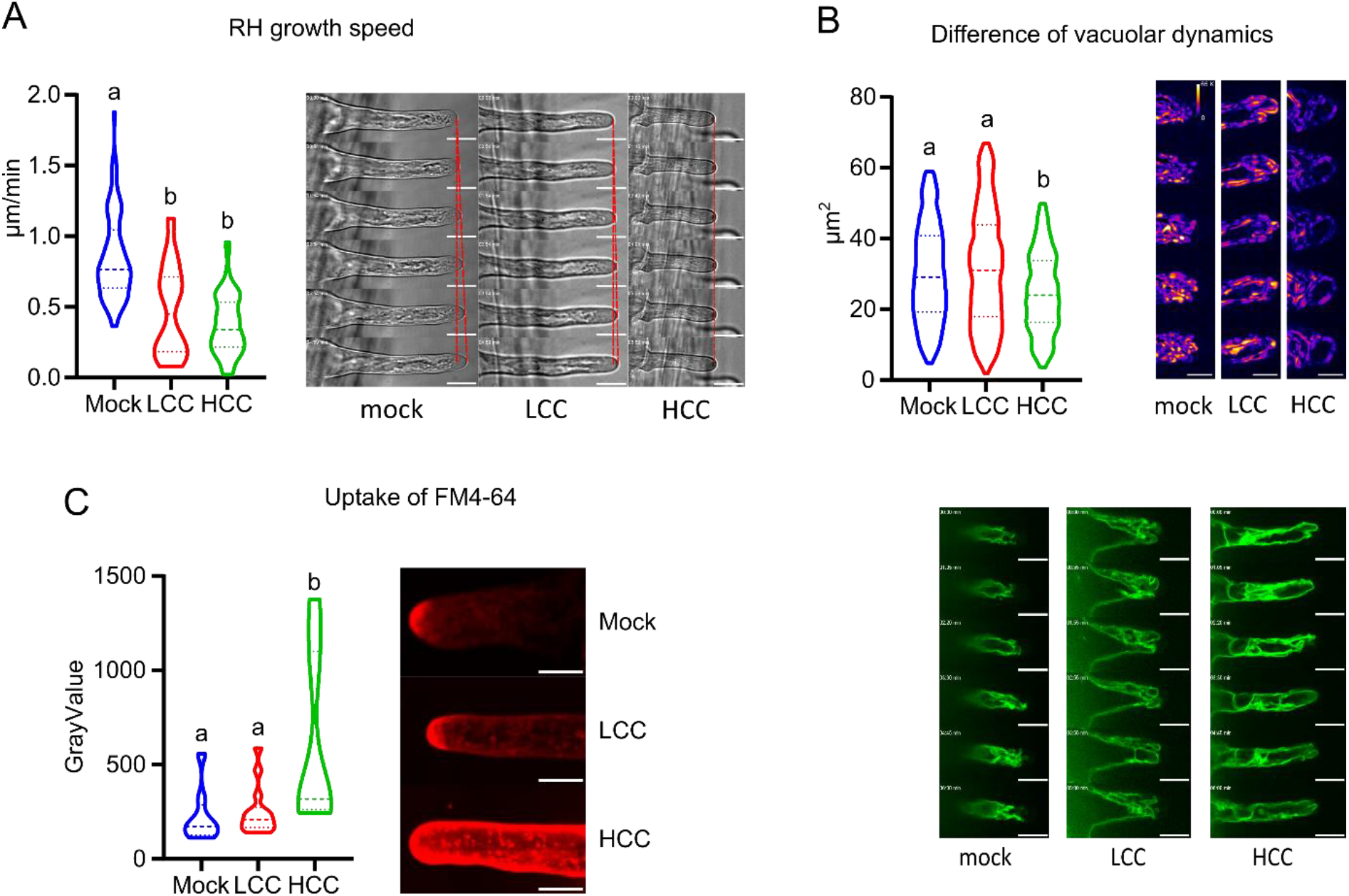
Subcellular dynamics during the early time points of LCC and HCC treatments. A) The real-time analysis of the root hairs growth during the mock, LCC, and HCC treatments based on the bright field images. The plot shows the RH growth rates for the mock, LCC and HCC treatments; small letters indicate significant differences based on Kruskal - Wallis test. B) The real-time changes in the vacuolar dynamics quantified as a vacuole-occupied area in the RH tip, imaged using the tonoplast marker Vti-GFP. The plot displays the differences between two consecutive frames; the examples of the differential signal are shown on the right; examples of vacuolar dynamics are shown below. Small letters indicate significant differences based on the ANOVA test. Bar = 20 µm. C) The analyses of the endocytosed FM4-64 dye after 15 min of treatments for mock, LCC, and HCC. The endocytic compartment stained by FM4-64 in the RH tip area is prominently enhanced in case of the HCC treatments. Small letters indicate significant differences based on Kruskal -Wallis test. On the right are examples of maximal intensity projections of FM4 -64 stained root hairs. Bar = 10 µm.

Since vacuolar dynamics is often governed by the actin cytoskeleton, we tested the effect of various chitosan concentrations on the organization and dynamics of filamentous actin in RHs. Somewhat unexpectedly, there was no difference in the actin organisation during the early response to chitosan, as dynamics of actin-decorating fimbrin-GFP were indistinguishable among the mock, LCC, and HCC treated plants (Supplementary figure 2).

The microscopical observations thus confirmed that the LCC, despite provoking prominent changes in CW composition and RH growth inhibition in later time points, does not impose a prominent stress that would disturb endomembrane compartments in early time points. On the other hand, HCC treatment significantly inhibits RH growth, rapidly affects calcium gradient and oscillations in RH tip, blocks vacuolar dynamics and provokes accumulation of endocytotic/FM4-64 marked compartment.

### Chitosan induces changes in gene expression and cell wall components

To obtain a more general view on chitosan responses and to gain another insight into distinct plant reactions to LCC and HCC treatment, we performed the RNAseq analysis of liquid medium-grown Arabidopsis seedlings treated with mock, LCC or HCC for 24 h. The general responsiveness of plants to LCC is far less dramatic than what is observed on the level of RHs, as only 58 differentially expressed genes (DEGs) were identified for LCC, and 8507 for HCC, comprising participants of plant PTI responses; most of the DEGs found for LCC are also present in the HCC DEG gene set, while only 11 genes are specifically regulated by LCC (Table I; Supplementary table I). Out of these 11 DEGs, 10 genes are downregulated, and only one, AtLRX6 (Leucine-rich repeat-extensin 6; AT3G22800), upregulated and found to belong to leucine-rich repeat/extension-type proteins, reported previously to play a role in RH growth. We have also identified genes which were differentially expressed in LCC and HCC treatments - in contrast to HCC which upregulates their expression, LCC downregulates (bottom 5 genes in the Table I). The restricted number of DEGs provoked by LCC is in concordance with milder Ca^2+^ and MAPK responses activation and with limited morphological changes observed mainly on the level of newly growing RHs, implying that the RH callose encasement is an effective but limited and localised strategy for the plant to overcome mild biotic threats, and to proceed with undisturbed development.

**Table I.**
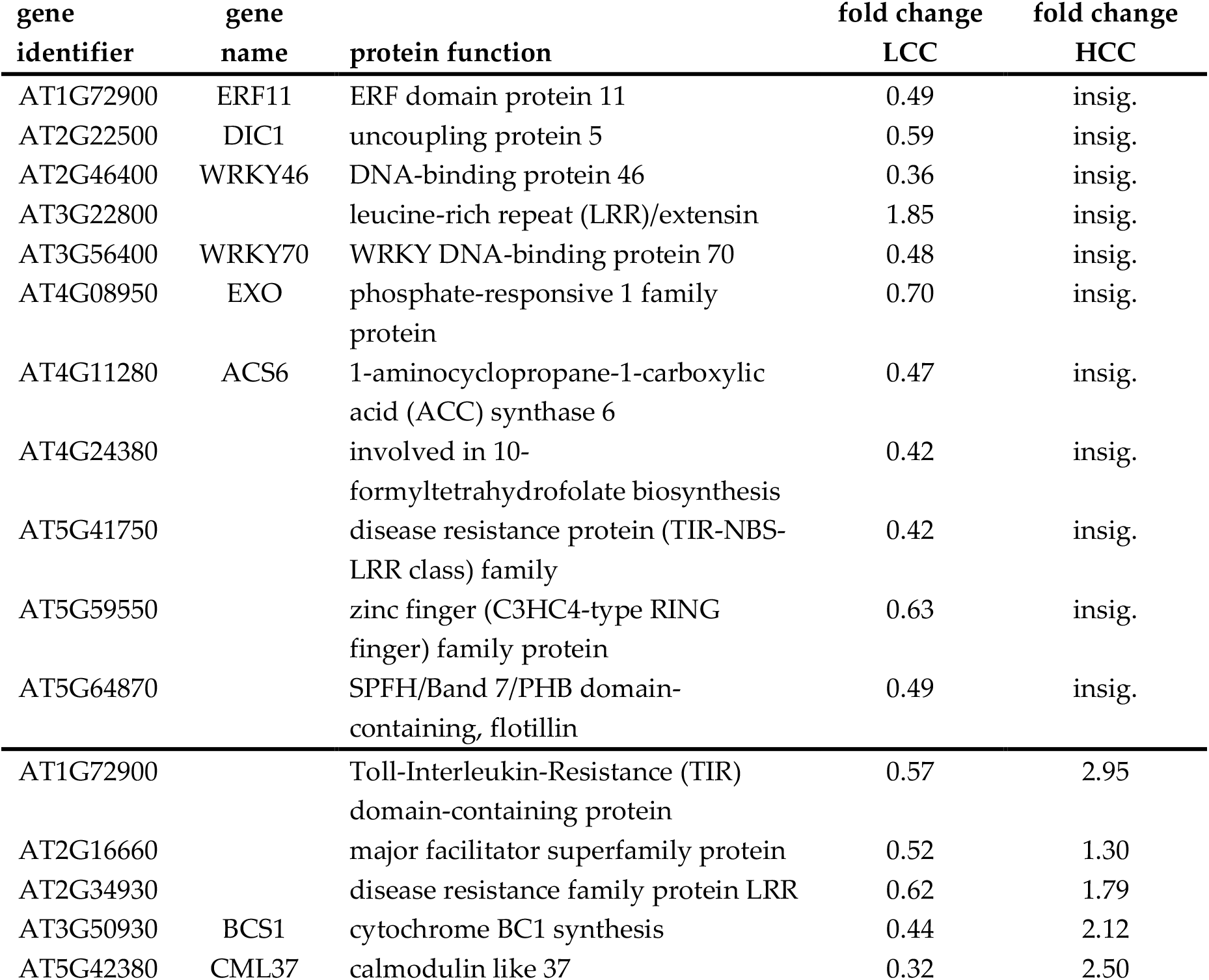
Eleven genes (out of 58 DEGS; most of them with high RH expression) are specifically affected by LCC, when compared to HCC (out of 8508 DEGs), as shown by the RNAseq analysis performed on 6 d.o. plants cultivated in liquid medium (mock, LCC, HCC; 24h); 5 genes below the line in the Table are differentially expressed in LCC in comparison to HCC (fold change < 1 = downregulation, fold change > 1 = upregulation).

Even though our results are centred around seedlings’ roots reactivity to chitosan, in order to obtain a more general view of the LCC impact on plants, we analysed LCC-caused gene expression of adult rosettes, and we encountered the results with those from the cell wall profiling of the LCC-treated rosette leaves (Supplemental figure 3A, 3B; (Voxeur et al., 2019)). Interestingly, the RNAseq of LCC-treated mature rosette leaves revealed differences that are more prominent and more similar to expression patterns provoked by other elicitor treatments, and mainly more similar to the HCC-treated seedlings. Both RNAseq and cell wall composition analyses accordingly revealed the affection of pectin synthesis and modification/methylation states upon LCC. We next realised an enzymatic fingerprinting of the homogalacturonans and xyloglucans using a commercial polygalacturonase and an endo-cellulase, respectively, from 5-weeks-old Arabidopsis leaves treated in presence of mock and LCC. We analysed the oligosaccharide produced by LC-MS (Paterlini et al., 2022). It revealed that LCC treatment likely triggered the demethylesterification and acetylesterification since the relative amounts of GalA4Me decreased and GalA3 Ac increased, respectively. Chitosan treatment also modified the structure and composition of hemicellulose building blocks. It remains to be further assessed whether, and how the CW components modifications take part in the RH responsiveness to chitosan.

### Callose deposition is conserved in evolutionarily unrelated structures functionally analogous to root hairs

Because the callose deposition was preserved in all tested signalling and/or secretory pathway mutants, we speculated that it is a robust reaction which might be evolutionary conserved and present also in non-angiosperm plant lineages and their stress responses. In a similar experimental setup with plants cultivated in the liquid medium, we were able to induce the growth of the morphological structures functionally analogous to root hairs, namely rhizoids in the liverwort *Marchantia polymorpha* and the rhizophores of spikemoss *Selaginella uncinata*. When the plants were treated for 24 h with LCC, *M. polymorpha* rhizoids growing out of an asexual reproductive body gemma were enclosed by callose, similarly to rhizophore surface cells of *S. uncinata* (Figure 7). Also, no callose depositions were detected in mock, while HCC treatment prevented their appearance, mirroring the situation with RH of *A. thaliana*.

**Figure 7.**
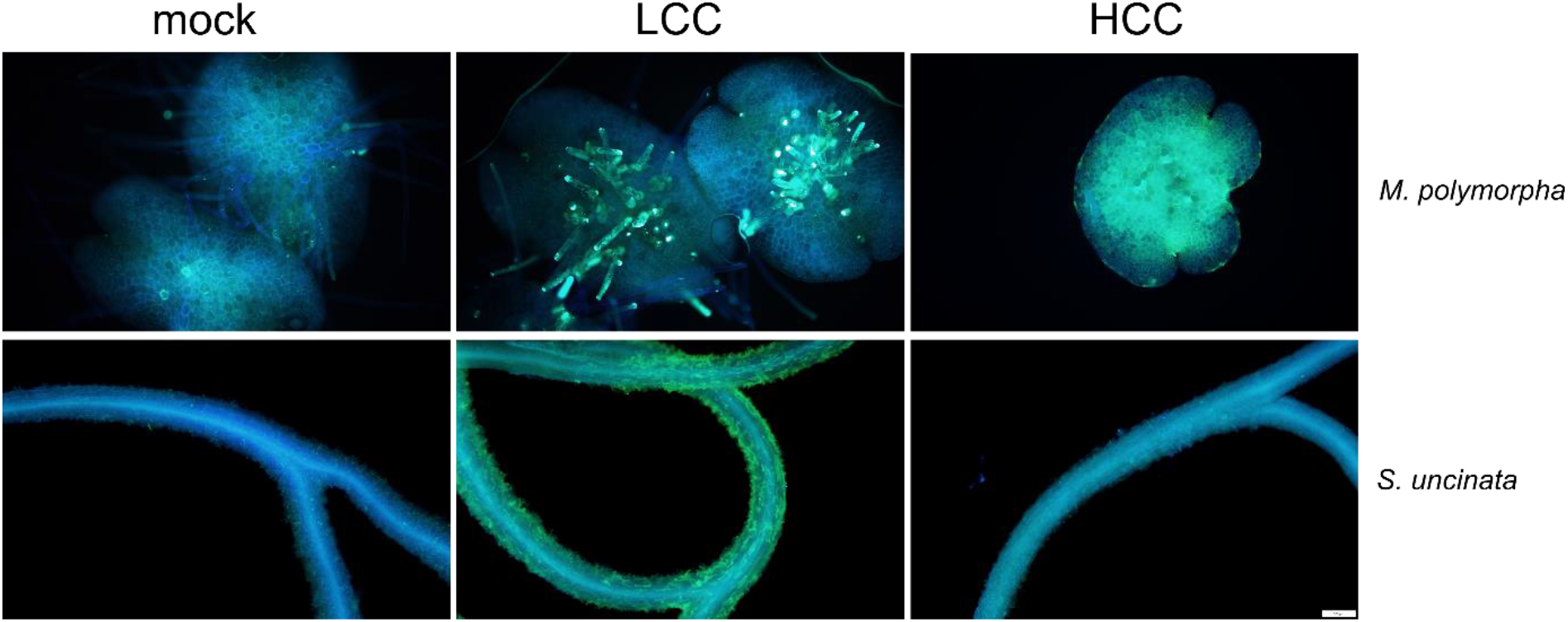
Callose deposition is conserved in structures analogous to root hairs. *M. polymorpha* (liverwort) rhizoids growing out of gemma are enclosed by callose when incubated for 24 h with LCC. The same effect is observable also on spikemoss rhizophores (*S. uncinata*). Bar = 100 µm.

We further verified if the relation of chitosan treatment and callose response can also be established for different unicellular model – regenerating protoplasts derived from the moss *P. patens* tip growing protonemal cells (Brejšková et al., 2021; Cove et al., 2009). We could indeed observe a prominent callose deposition caused by LCC, while HCC treatment had a deleterious effect on protoplast viability and regeneration (Supplementary figure 4A). Interestingly, in contrast to *P. patens* regenerating protoplasts, above-ground tissue-derived protoplasts of *A. thaliana* did not deposit callose while regenerating in the presence of chitosan (Supplementary figure 4B).

These results confirmed that the potential of RH and RH-like structure to deposit callose is an evolutionary old intrinsic feature. This capacity, however, may be exclusive for the tissue derived from the non-aerial plant parts, exhibiting fast cellular growth and newly generating the cell wall.

## Discussion

In this work, we uncovered the RHs capability to perceive the chitosan as an elicitor, and depending on its concentration, to deposit callose to the cell wall and/or to inhibit RH growth. The two processes are not mutually dependent, and the two studied chitosan concentrations, the low (LCC) and the high ones (HCC), may induce plant responses that overlap in their quality and quantity, but also those which are specific. We also demonstrate that the shift in treatment stringency toward the milder concentrations of the stressor can reveal new defence- and development-related phenotypes.

Phytopathological studies often deal with approaches involving high elicitor concentrations necessary to provoke responses that could be easily observed and quantified (Aslam et al., 2009; Luna et al., 2011). We speculate that the low chitosan concentration (LCC, 0.001%), used in our study, imposes a mild biotic stress, which young plants can sustain, and to which they react by the adjustments on the level of RHs (as the most exposed cell type) by slowing down their growth rate and by fortifying the cell wall with callose. Besides RHs, the LCC stimuli are also perceived by other root parts and also above-ground organs, as our results from Ca^2+^ and MAPK assays showed. Interestingly, prolonged LCC treatment had no impact on the growth of young plants, and they are not affected in their further development, implying an accommodation to these low chitosan concentrations, also observable in quenching of early phases of Ca^2+^ spiking patterns. In agreement with early signalling patterns, and results of the RNAseq and CW analysis, the susceptibility assays showed an enhancement of resistance of plants to Pst *hrcC* bacteria when applied upon HCC pretreatment, while the effects of LCC leaves no durable effect on the immunity of young seedlings.

Similarly, the RNAseq data from LCC- and HCC-treated seedlings corroborate the existence of primarily quantitative differences between the two treatments since only a minority of DEGs was found to be specific for LCC. Despite the scarcity of the LCC-specific candidates, we consider that the role of one of the identified DEGs, LRX10, is worthy of further verification, since this protein, together with related pollen tube-expressed isoforms, has been found to be involved in the pollen tube cell wall pectins and callose deposition, important for the process of fertilisation (Wang et al., 2018) Therefore, the presence of minor qualitative differences is legible as well. This is further fortified by the fact that, unlike the HCC treatment, which impairs plant growth by affecting the auxin pathway (Lopez-Moya et al., 2017), the LCC has only a minor effect on auxin signalling enhancement, as demonstrated using the DR5::GFP auxin reporter line (Supplemental figure 5).

Callose deposition is typically described as an early defence response in leaf tissue, where it serves mainly to seal the plasmodesmatal connection between cells to prevent pathogen spread and nutrient withdrawal (Epel, 2009) or to prevent pathogens’ entrance (Meyer et al., 2009; Ortmannová et al., 2022)), but also in primary root epidermis after the defence-eliciting treatments, displaying a similar spotty pattern as found in leaves (Millet et al., 2010). In agreement with this defence involvement, callose deposition is repressed in the establishment of mutualistic interactions (rev. (German et al., 2023)). Besides the defence-related functions, callose also serves as a mechanical support in the new cell wall formation, in the pollen exine formation and germinating-pollen plug, but also in a functional specialisation of trichome basal parts (Kubátová et al., 2019; Kulich et al., 2015; Ma et al., 2021; Motomura et al., 2022). The RH callose deposition observed by us is tightly coupled with a state of the elongating root cells, and in the newly growing root hairs in the zones closest to the root tip. Therefore, we hypothesise that the RH callose coating is the consequence of two processes going along – of the elicitor caused defence signalling, as evidenced by the dependance on the typically pathogen-induced isoform of callose synthase PMR4, but also of the fragile and immature state of the cell wall undergoing synthesis, which was additionally confirmed on our experiments with regenerating protoplasts that readily deposit callose upon the LCC treatment (Dehors et al., 2019), as well on recently demonstrated abiotic stress and starvation related RH callose deposition (Okada et al., 2023).

Both LCC / HCC treated rhizoids/rhizophores of *M. polymorpha* and *S. uncinata*, despite their evolutionary unrelated origins, provide the same morphological and subcellular responses, including the employment of the evolutionarily conserved callose synthases. This functional conservation in terrestrial plants might have an origin in the callose contribution to the newly formed post-meiotic cell wall of spores or pollen. Secondarily, these independently-evolved mechanisms may have also been involved in the onset of responses of plant cells to the hostile environmental conditions of land/soil, also including pathogen threat (Herburger and Holzinger, 2015; Ks et al., 2020).

Unlike LCC, the HCC has a prominent effect on the appearance and subsequent loss of complexity of the RH tip-localised portion of the vacuolar compartment. The HCC causes the vacuolar withdrawal and dismission of the complex tubular tip-localised vacuolar growing parts. Given the massive uptake of FM4-64 dye in HCC-treated RHs, this may be the consequence of the enlargement of the endocytic zone in the RH tip, or an unrelated endomembrane response. The prompt response of vacuolar dynamics to chitosan may suggest that, besides a proposed role for vacuolar calcium channels in mechanosensing of the tip-growing cells, a part of the pathogen sensing might reside on the tonoplast as well (Radin et al., 2021).

Despite the critical importance of the actin cytoskeleton in both RH growth and responsiveness to pathogens, in our experiment, at an early time points, there was no significant modification of fimbrin-GFP-labelled actin filaments upon chitosan treatments (Supplementary figure 2, see also (Porter and Day, 2016; Ringli et al., 2002; Sun et al., 2021; Vaškebová et al., 2018)). It would be interesting to verify if this observation might be related to the different requirements for the callose- and cellulose synthases localization along the plasma membrane by the microtubules and the actin, respectively, similarly to what has been found for the tobacco pollen tube (Cai et al., 2011).

In addition to demonstrating the usefulness of mild chemical/eliciting treatments of plants for opening new perspectives in plant studies, we are also suggesting the simple experimental setup based on a single, exposed, and sensitive cell. This approach could be further employed for gaining the knowledge on the root and RH-related immunity, as well as in the studies of the overall plant fitness in relation to the rhizosphere conditions.

## Conclusions

The effect of the low concentration chitosan treatment on Arabidopsis root hairs, and similar effect in Marchantia and Selaginella rhizoids or rhizophores, expose their unexpected deeply rooted capacity to react by the callose accumulation in response to this fungal elicitor. The root hair cell wall callose deposition is PMR4/GSL5-dependent and occurs in parallel but independently of the root hair growth inhibition. The higher chitosan concentration treatments cause prominent root hair growth inhibition that precludes the callose deposition, leading to the phenotypic difference between the two treatments. The intensive root hair growth inhibition is probably the consequence of the emphatic defence alerting, as well as interference with the root tip-localised vacuolar and endocytic compartments.

## Material and methods

### Plant cultivation

For seedlings and plants cultivation, seeds were surface sterilised (5 min in 70% ethanol, 2 × 5 min in 10% commercial bleach, rinsed three times in sterile distilled water) and stratified for 2–3 days at 4°C. Seeds were then germinated and grown on vertical ½ MS agar plates (half-strength Murashige and Skoog salts, Duchefa Biochemie, supplemented with 1% sucrose, vitamin mixture, and 1.6% plant agar, Duchefa Biochemie) at 21 °C and 16 h of light per day for five-seven days. For propagation of plants, seedlings were transferred into Jiffy Products International pellets and grown at 22 °C and 16 h of light per day in growth rooms.

The following previously published Arabidopsis mutant and transgenic lines were used in this study: *aux1-21* (CS9584, (Bennett et al., 1996)), *bak1-4* (CS71777; (Kemmerling et al., 2007)), *cals8-1* (SALK_037603; (Cui and Lee, 2016)), *cerk1-1* (pst14772; (Miya et al., 2007)), *cngc14-1* (SALK206460; (Shih et al., 2015)), *cngc2-3* (SALK_066908; Chin et al., 2013), *exo70H4-1* (SALK_023593; (Kulich et al., 2015)), *npr1* (CS3726; (Cao et al., 1994)), *pmr4-1* (CS67159; (Vogel and Somerville, 2000)), *rbohD-3* (N9555, Torres et al., 2002 11756663), UQ10::GFP-PMR4 (Kulich et al., 2018), promPMR4::GFP-PMR4 (Sabol et al, in preparation), DR5-GFP (Heisler et al., 2005), fimbrin-GFP (Wang et al., 2004), and vacuolar ATPase Vti-GFP (gifted by Karen Schumacher, Heidelberg). The wild type (WT) Col-0 was used as control. When needed, primers used for the genotyping were the same as used in the above-cited original studies.

### Chitosan treatment

Plants were propagated in vitro for 5-7 d on vertical plates, and then transferred by forceps into liquid ½ MS (control) or liquid ½ MS with chitosan added (100x and 1000x diluted stock solution of the 1% chitosan in 1% acetic acid), usually 4-8 plants per 3 ml, in 6-multiwell plates (Nunclon, Thermo Fisher Scientific). Plants were cultivated for 24 h 16/8 h light/dark at 21–22°C without shaking. After 24 h of incubation, plants were transferred to 3 ml of ethanol for 6-24 h at room temperature for destaining, with mild shaking, and subsequently transferred into 3 ml of 150 mM K2HPO4. After 1 h, aniline blue (final concentration 0.005% solution in 150 mM K2HPO4) was added and seedlings stained for additional 24 h with mild shaking and observed under the microscope. Additional verifications confirmed that there were no differences between the mock treatment with and without corresponding dilution of acetic acid. Several chitosans used for analyses were purchased from Merck, Germany (Cat. Nos C3646, 523682, 448877 and 419419), as well as chitin (C7170).

### Root and root hair phenotype examination

The plant responses to pathogens were monitored by microscopy and documented by photography. The seedlings were placed on microscopic glass, their primary root stretched and lengths measured. Microscopic analysis of root tips and root hairs was performed using an Olympus BX51 microscope with attached DP50 camera (Olympus). For each treatment/genotype 5-10 plants were photographed and analysed. Root hair lengths were determined for the first 20-40 root hairs starting from the root tip for each plant; for statistical analysis. Callose deposition was analysed using the pre-set conditions for bright-field and DAPI, and objective 5x or 10x.

### Protoplasts preparation

The moss *Physcomitrium patens* Gransden strain was routinely propagated in vitro on BCD medium supplemented with ammonium tartrate dibasic (BCDAT), according to (Cove et al., 2009), in a climate chamber (16 h of light/8 h of dark, 25 °C, illuminated by fluorescent tubes at 50-70 µmol m^-2^ sec^-1^). Protonemal tissue (6-7 days after propagation) from cellophane-grown culture was transferred into the 1%Driselase/8%mannitol solution. When filamentous structure became invisible (approx. 1 h at RT), protoplast suspension was filtered via/through 100 µm steel mesh. Protoplasts were diluted in 8% mannitol up to 10 ml and spun down (700 rpm for 4 min, without brake) and this step was repeated three times. Protoplast sediment was carefully resuspended between centrifugations to avoid aggregation of protoplasts. Finally protoplasts were incubated in the liquid regeneration medium (BCDAT+ 5mM NH_4_+ 6% w/v mannitol + 10 mM CaCl_2_) overnight at RT. Arabidopsis protoplasts were prepared according to (Yoo et al., 2007). Briefly, Arabidopsis leaves from 3-4 weeks old plants grown under short day conditions (8/16 h light/dark) were cut onto 2 mm stripes. The 2 g of prepared plant tissue were submerged in 37 °C warm protoplast solution (0.4M mannitol, 1.5% cellulase, 0.4% macerozyme, 20mM MES pH 5.7, 20mM KCl, 0.1% BSA, 10mM CaCl2) and vacuum has been applied for 30 min. The submerged leaves were kept in the dark in RT. After 3 h, the flasks with protoplasts were gently swirled and the quality of protoplasts was checked with a microscope. The protoplast solution was washed with a WI buffer (4mM MES pH 5.7, 0.5M mannitol, 20mM KCl), filtered through miracloth and clarified with multiple centrifugations (3x100g, 2 min). For the recovery of protoplasts, 0.7 ml of WI solution in a 12-well plate was applied. Both types of protoplasts were left to recover for 3h, and were subsequently treated with chitosan or mock for 24 h. Protoplasts were stained with aniline blue and the final condition of protoplasts monitored with the microscope Olympus BX51 microscope with attached DP50 camera.

### Selaginella and Marchantia propagation and callose deposition assay

For sterile *Selaginella uncinata* (‘Comenius’ strain) culture establishment, excised fragments containing meristems were washed with distilled H_2_O and surface sterilized with 70% EtOH (1 minute) and subsequently with 40% commercial bleach (SAVO)/2% sodium hypochlorite for 15 minutes and rinsed with a sterile water twice (Park et al., 2020). Axenic regenerants were subsequently vegetatively propagated on ½ MS media with 0.6% agar under standard conditions (22°C, 16/8 light/dark, ca. 180 µmol/m^2^/s). Chitosan treatments were performed on whole regenerating plantlets interchangeably with the *Arabidopsis thaliana* approach.

*Marchantia polymorpha* Tak-1 has been maintained on ½ Gamborg media ((Gamborg et al., 1968); G0210 Duchefa) supplemented with 1,2% agar under standard conditions (22°C, 16/8 light/dark, ca. 180 µmol/m^2^/s). Chitosan treatments were done with gemmae analogously to an *Arabidopsis thaliana* procedure, with a slight change, where the liquid ½ MS has been replaced with ½ Gamborg media supplemented with 1% sucrose to enhance the rhizoids initiation.

### Confocal and Real Time microscopy

Small agar block with *in vitro* cultivated 6-day-old seedlings was gently cut from the cultivation plate and transferred into sterile microscopic chambers containing 100 ul of liquid ½ MS. Chambers were covered with lids and left overnight in a cultivation chamber. For real time treatments we used custom designed 3D printed perfusion add-on which fits into the microscopic chamber and allows exchange of media inside the microscopic chamber during microscopy session. To mediate the exchange we used medical grade silicone tubes mounted onto a peristaltic pump and perfusion add-on with speed approximately 100 ul/min. All microscopic samples were 10 min pretreated with the mock solution right before capturing, chitosan treatment was applied directly during the microscopic session. For mock treatments the liquid 1⁄2 MS was used. For chitosan treatments, 0.001% chitosan (LCC) or 0.01% chitosan for HCC were added into the ½ MS. For the FM4-64 (Thermofisher scientific) staining we used the same perfusion settings and solutions just with the addition of 0.001% FM4-64 dye (1 mM stock solution). Samples were treated for 15 min before observation.

For RH R-GECO1 signal intensity analyses the same microscopic perfusion add-on was used. The 5-6 day old RGECO1 expressing seedlings were transferred into microscopic chambers, covered with a lid and kept overnight in the cultivation room. Samples were treated using the same chitosan solutions as in previous experiments (LCC, HCC). Roots were captured every 1 second for 7 min for each part (untreated, mock, LCC, HCC) in the case of LCC and HCC we added 30 seconds long (capturing time one frame per 0,1 second) to monitor fast changes. These super-fast scanning parts were removed for the figures.

Real Time imaging was done on the Spinning disc (SD) microscope Nikon (Eclipse Ti-E, inverted) with Yokogawa CSU-W1 SD unit (50mm) equipped with Omicron LightHUB ULTRA light source, Plan-Apochromat L 20x/0.75 or Plan-Apochromat LS 40x/1.25 WI objectives were used for capturing. Microscope is operated by NIS Elements 5.30 software. GFP was observed using a 488 nm excitation laser with Semrock brightline Em 525/30 filter. FM4-64 was observed using a 488 nm excitation laser with Semrock brightline Em 641/75 filter.

**F**or the analysis of vacuolar dynamics a *stack difference* task inside *mutli kymograph* plugin ImageJ (FIJI) followed by analysis of *moments* thresholded area of the result image was used. See Supplementary material *VD_macro.ijm*.

For the FM4-64 uptake analysis the average Z projections of single RH were taken with subsequent analysis of signal intensity within inner cell space (ImageJ - FIJI). Actin dynamics wes analysed using the same procedure but with longer gap between consecutive frames (10 seconds).

### Ca2+ signalling reporter imaging and evaluation

*A. thaliana* seedlings expressing R-GECO1 Ca2+ signalling reporter were vertically grown in a growth chamber at 23 °C by day (16 h), 18 °C by night (8 h), 60% humidity and light intensity of 100 µmol photons m^−2^ s^−1^ (Keinath et al., 2015). Microfluidics experiments were performed using four-day-old seedlings placed into a microfluidic chip and enclosed by a cover glass (Serre et al., 2021). Channel one contained a control solution (½ MS, fluorescein dextran 6 µg/ml), while channel two contained a treatment solution (½ MS, 0.01% or 0.001% chitosan). The media flow rate was 3 µl/min (OBI1, MFS2 Elveflow, and Elveflow software ESI (v.3.04.1). The system was mounted to a vertical microscope stage and kept for 15 min plant recovery. The media exchange was indicated by fluorescein-dextran signal intensity.

Plants were imaged using a vertical stage Zeiss Axio Observer 7 with a Yokogawa CSU-W1-T2 spinning disk unit with 50-µm pinholes equipped with a VS-HOM1000 excitation light homogenizer (Visitron Systems), objective Zeiss Plan-Apochromat ×10/0.45. Images were acquired using VisiView software (Visitron Systems, v.4.4.0.14). R-GECO1 was excited by a 561 nm laser, and fluorescein dextran by a 488 nm laser. The signal was detected using a PRIME-95B Back-Illuminated sCMOS camera (1,200 × 1,200 pixels; Photometrics). The seedlings were imaged every 13 s for 60 min. The signal intensity of R-GECO1 was measured inside the root at the root late elongation zone using ImageJ Fiji software (Schindelin et al., 2012). The intensity of the background was measured in areas not affected by the root response and subtracted from the R-GECO1 values. The intensity of fluorescein-dextran was measured outside of the root. The data were normalised using division by initial fluorescence intensity values (F/F0). As the initial value F0, we used the average fluorescence intensity at five-time points before treatment.

### RNA isolation and RNAseq analysis

RNA was isolated from young, 6-7-day-old whole seedlings (transferred from vertical ½ MS plates to liquid ½ MS for treatments), and adult (5-6 week old) rosette leaves of *A. thaliana* plants, using the RNeasy Plant kit (Qiagen) according to the manufacturer’s instructions. Isolated RNAs were stabilised by GenTegra technology microtubes (GenTegra, Pleasanton, California, USA). Strand-specific cDNA libraries were constructed from polyA enriched RNA and sequenced on the Illumina NovaSeq6000 platform with subsequent analysis performed by Eurofins. The RNA-seq data used in this study are deposited in the National Center for Biotechnology Information Gene Expression Omnibus database under the accession number GSE239633.

### Protein extraction, western blot and MAPK assay

Total protein extracts were isolated from 5–7-day-old *A. thaliana* seedlings according to procedure described in (Fernandez and Beeckman, 2020). Briefly, untreated and chitosan treated tissue samples were ground in liquid nitrogen and dissolved in an extraction buffer containing Phosstop (Roche), and the concentration of proteins determined by the Bio-Rad protein assay kit with bovine serum albumin (BSA) as the standard. The extracts were denatured by boiling in a 6× SDS loading buffer. The protein samples were separated by 10% SDS-PAGE and analysed by Western blot using the α-p44/42-ERK antibody (SAB4301578), anti-MAPK3 (M8318, Sigma), anti-MAPK4 (A6979, Sigma), and anti-MAPK6 (A7104, Sigma). The primary antibodies were incubated with the membranes for 3 h at room temperature in the blocking solution. Horseradish peroxidase-conjugated antibodies (anti-rabbit and anti-mouse; Promega) were applied followed by chemiluminescent ECL detection (Amersham) by the Bio-Rad documentation system. Using the Gel Analysis function of ImageJ, signal intensities for protein bands were determined for each treatment from 3 different samples. Loading consistency was examined by staining the membrane with Ponceau S.

### Bacterial assay

Bacteria susceptibility assay was performed according to Ishiga et al., 2011 and Pečenková et al., 2020 with modifications. Briefly, plants grown in vitro (½ MS, at 21°C, 12 / 12 h of light / dark per day, for 6 days) were transferred to liquid ½ MS in 6-well plates, and pretreated with mock, or with chitosan (LCC and HCC). After 24 h, the medium was exchanged, and the *P. syringae pv. tomato hrcC* (Pst *hrcC*) OD = 0.01, was used for seedlings inoculation. After the 24 h incubation with mild shaking 50 rpm/min, ten seedlings in one sample, always three samples for each treatment, in two replications) were homogenised, homogenates subsequently diluted in series, plated out, and after approximately 24-36 h of incubation on 28°C colonies counted.

### Cell wall preparation and enzymatic fingerprinting

For cell wall analysis plants have been propagated in Jiffy pellets for 5-6 weeks, 10/14 h of light/dark regime. Plants were treated by dipping into the mock (distilled water with 0.005% Silwett) or LCC chitosan (distilled water with 0.005% Silwett and 0.001% chitosan), and left for 24 h in the growth chamber. Treated and non-treated samples were submerged in 96% (v/v) ethanol and boiled at 70°C for 10 min. The pellets were collected by centrifugation (13000 g for 10 min) and dried in a speed vacuum concentrator at 30°C overnight. Samples were digested with 1 U/mg DW sample of *Aspergillus aculeatus* endo-polygalacturonase M2 (Megazyme, Bray, Ireland) (Paterlini et al., 2022) in 50 mM ammonium acetate buffer (pH 5) at 37 °C for 18 h. Samples were centrifuged at 13000 rpm for 10 min and 100 µL of the supernatants were transferred into vials. The oligosaccharides released from digestion were analysed according to (Paterlini et al., 2022).

### Image analysis and statistics

Image processing software was employed for image data quantification. Roots and root hair sizes were analysed using AnalySIS (Soft Imaging System GmbH, Germany) or ImageJ (Schneider et al., 2012) software. The numerical data obtained were processed using Microsoft Excel.

To determine statistical significance, Student’s t-test and ANOVA tests were conducted either using Excel or on-line calculators (http://in-silico.net, and http://statpages.info/anova1sm.html). P-values of less than 0·05 indicated a significant difference among the various groups. For multiple-comparison experiments, the Kruskal - Wallis or Tukey Honestly Significant Difference (HSD) post hoc test were used.

## Contribution

M.D. performed root hair confocal microscopy, designed the experimental setup for the real time observation of root hairs, performed microscopy and image analysis, prepared RNA for RNAseq analysis, prepared figures and wrote the manuscript. S. H. performed Selaginella and Marchantia experiments, and wrote the corresponding parts in the manuscript. E. Š. performed MAP kinases assays. P. K. and M. F. performed Ca2+-spiking recording and analysis. A. P. performed Pst *hrcC* susceptibility assays. L. B. and J. O. prepared protoplasts and analysed their cell wall. K. M. and N. S. analysed RNAseq data. A. V. and S. V. did the cell wall fingerprinting analysis. G. C. verified callose deposition in mutant lines. M. P. and V. Ž. planned the experiments and wrote the manuscript. T. P. performed root hair growth and callose deposition analysis, prepared figures and wrote the manuscript.

## Acknowledgment

We would like to thank to Norbert Zlámal from the Bratislavàs Comenius University Botanical Garden for sharing *Selaginella uncinata*, Melanie Krebs (Uni Heidelberg), Adriana Jelínková and Edita Janková Drdová (IEB CAS) for sharing Arabidopsis mutants and markers seeds, and Jana Šťovíčková for technical assistance. This work was supported by the Ministry of Education, Youth and Sports (MEYS) project CZ.02.1.01/0.0/0.0/16_019/0000738, 8J19FR001 MEYS Czech-French mobility program and Charles University Grant Agency (GAUK) project 415322. The Imaging Facility of the Institute of Experimental Botany AS CR is supported by the MEYS CR (LM2023050 Czech-BioImaging) and IEB AS CR.

---

**Supplementary** figure 1. A) Cultivation in water does not favour the RH growth and callose deposition; the RH callose reaction requires a rich cultivation medium in order for RHs to grow, and callose deposits to be formed. B) The growth of new RH is inhibited in early time points (4 and 8 h p. t.) in high chitosan concentration (HCC; bars = 100 µm). C) Arabidopsis mutants in secretion, defence and phytohormone signalling have ability to deposit root hair callose similarly to WT/Col-0. Bars = 100 µm.

**Supplementary** figure 2. Actin dynamics is not significantly altered during the early time points of the HCC treatment, as observed using fimbrin-GFP marker and quantified using the variance between the two time points during the observation; images on the right show examples of 5 consecutive frames for the two treatments.

**Supplementary** figure 3**. A)** Numbers of **s**pecific and common DEGs among three different types of experiments with chitosan treatments. B) Cell wall changes triggered by LCC treatment. Enzymatic fingerprinting of homogalacturonans (upper graph) and xyloglucans (lower graph) from cell walls of WT mock and LCC treated leaves. Data represents mean ±SD, n=4, *P<0.05, **P<0.01, student’s t test. OGs are named GalAxMeyAcz. x, y and z indicate the DP and the number of methyl- and acetylester groups, respectively. GalA: Galacturonic acid, Me: methylester group, Ac: Acetylester group. Xyloglucan fragment nomenclature follows that described in Fry et al. (1993): G stands for glucose; X for xylose-glucose; L for galactose-xylose-glucose; F for fucose-galactose-xylose-glucose.

**Supplementary** figure 4. Chitosan LCC treatment induces deposition of callose in moss *P. patens* regenerating protoplasts (A) but not in leaf mesophyll-derived Arabidopsis regenerating protoplasts (B), as assayed by aniline blue staining. Bars = 100 µm.

**Supplemental** figure 5 LCC has a minor effect on auxin signalling enhancement as demonstrated using the DR5::GFP reporter line (bar = 100 µm).

