## Supplementary Figure for "Chitosan stimulates root hair callose deposition and inhibits root hair growth"

Drs et al.

**Supplementary figure 1.** A) Cultivation in water does not favour the RH growth and callose deposition; the RH callose reaction requires a rich cultivation medium in order for RHs to grow, and callose deposits to be formed. B) The growth of new RH is inhibited in early time points (4 and 8 h p. t.) in high chitosan concentration (HCC; bars = 100 μm). C) Arabidopsis mutants in secretion, defence and phytohormone signalling have ability to deposit root hair callose similarly to WT/Col-0. Bars = 100 μm.

**Supplementary figure 2**. Actin dynamics is not significantly altered during the early time points of the HCC treatment, as observed using fimbrin-GFP marker and quantified using the variance between the two time points during the observation; images on the right show examples of 5 consecutive frames for the two treatments.

**Supplementary figure 3. A)** Numbers of specific and common DEGs among three different types of experiments with chitosan treatments. B) Cell wall changes triggered by LCC treatment. Enzymatic fingerprinting of homogalacturonans (upper graph) and xyloglucans (lower graph) from cell walls of WT mock and LCC treated leaves. Data represents mean ±SD, n=4, *P<0.05, **P<0.01, student's t test. OGs are named GalAxMeyAcz. x, y and z indicate the DP and the number of methyl- and acetylester groups, respectively. GalA: Galacturonic acid, Me: methylester group, Ac: Acetylester group. Xyloglucan fragment nomenclature follows that described in Fry et al. (1993): G stands for glucose; X for xylose-glucose; L for galactose-xylose-glucose; F for fucose-galactose-xylose-glucose.

**Supplementary figure 4.** Chitosan LCC treatment induces deposition of callose in moss *P. patens* regenerating protoplasts (A) but not in leaf mesophyll-derived Arabidopsis regenerating protoplasts (B), as assayed by aniline blue staining. Bars = 100 μm.

**Supplemental figure 5** LCC has a minor effect on auxin signalling enhancement as demonstrated using the DR5::GFP reporter line (bar = 100 μm).

**Supplementary figure 1.**

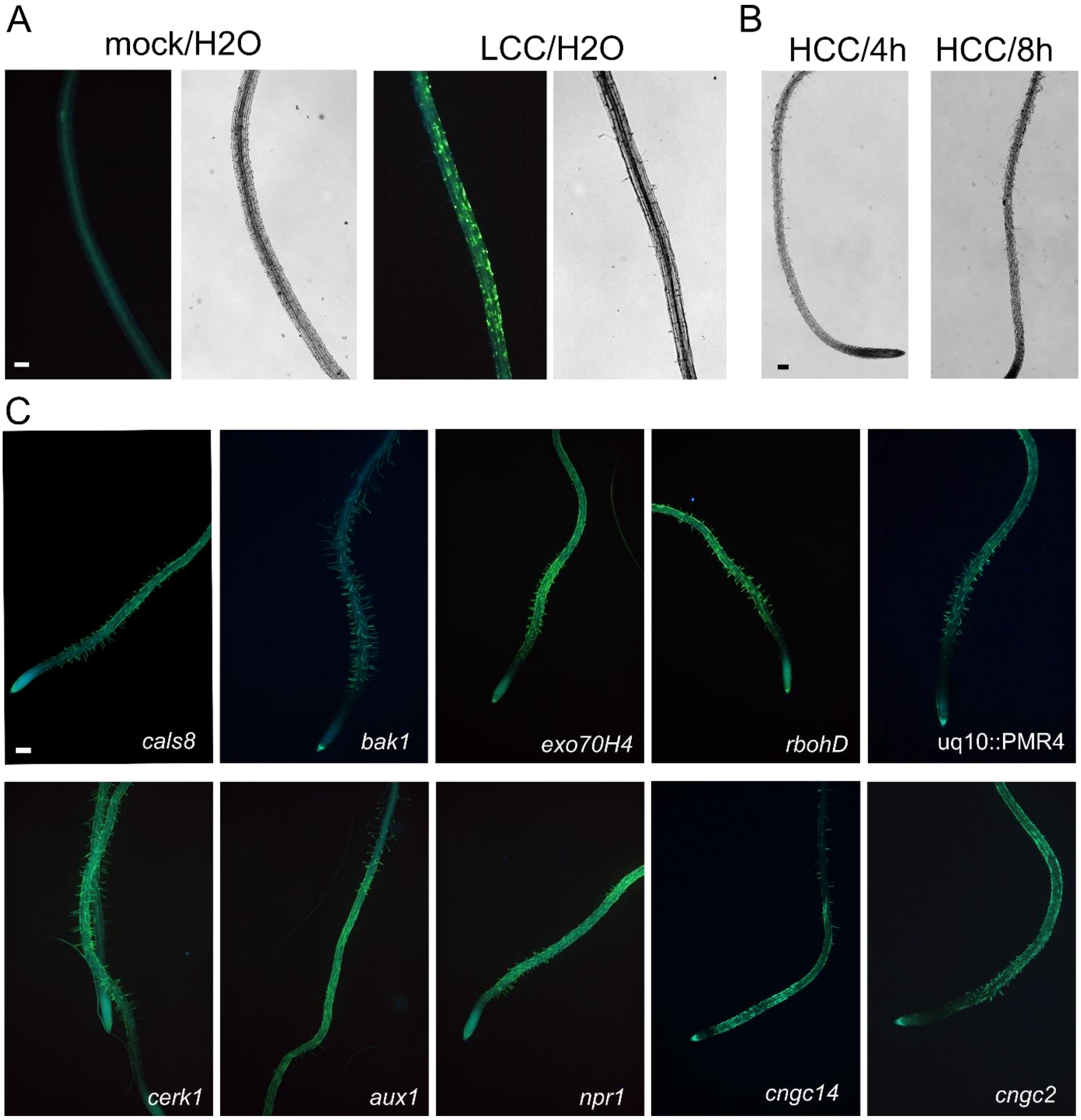

**Supplementary figure 2.**

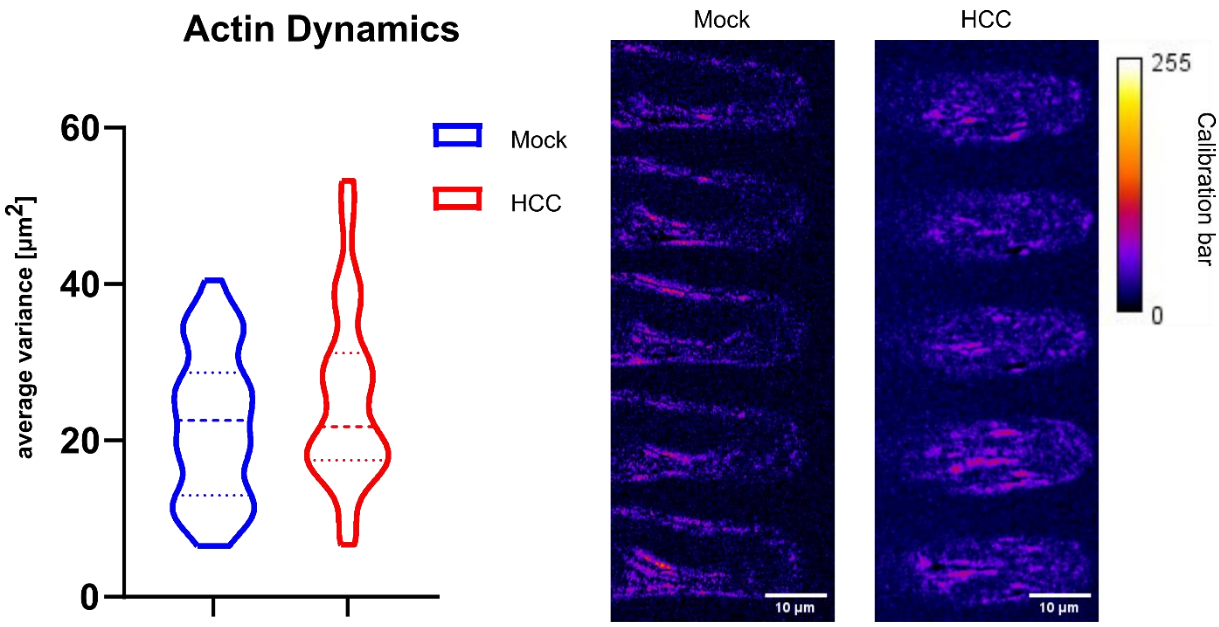


**Supplementary figure 3.**

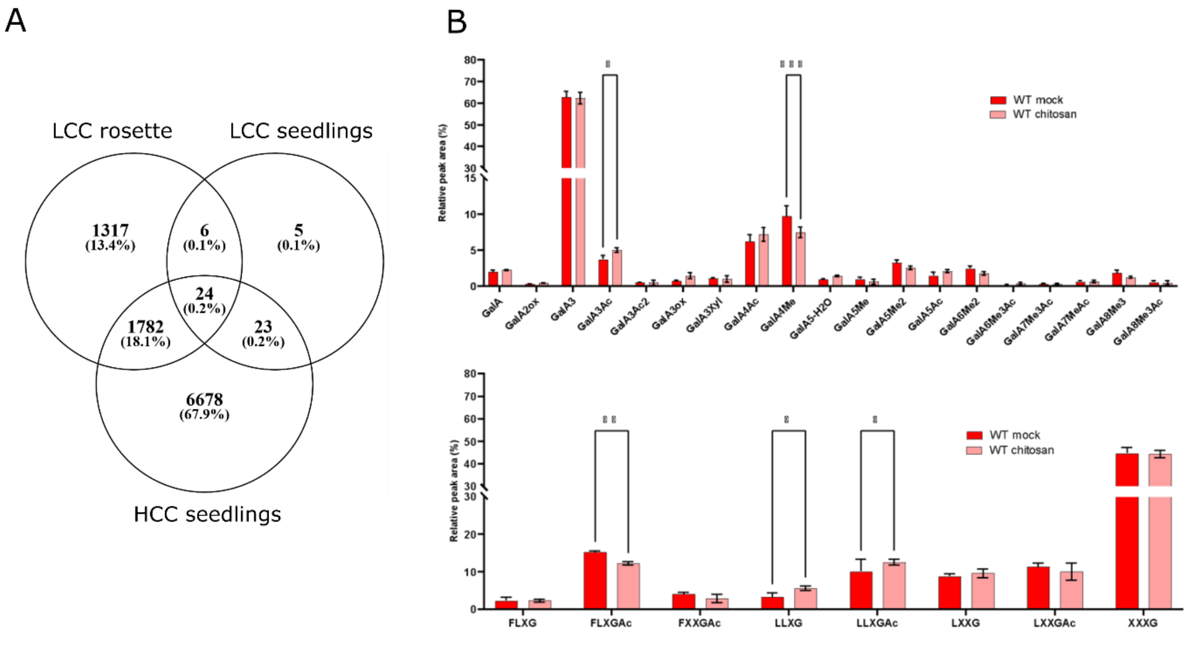

### Supplementary figure 4.

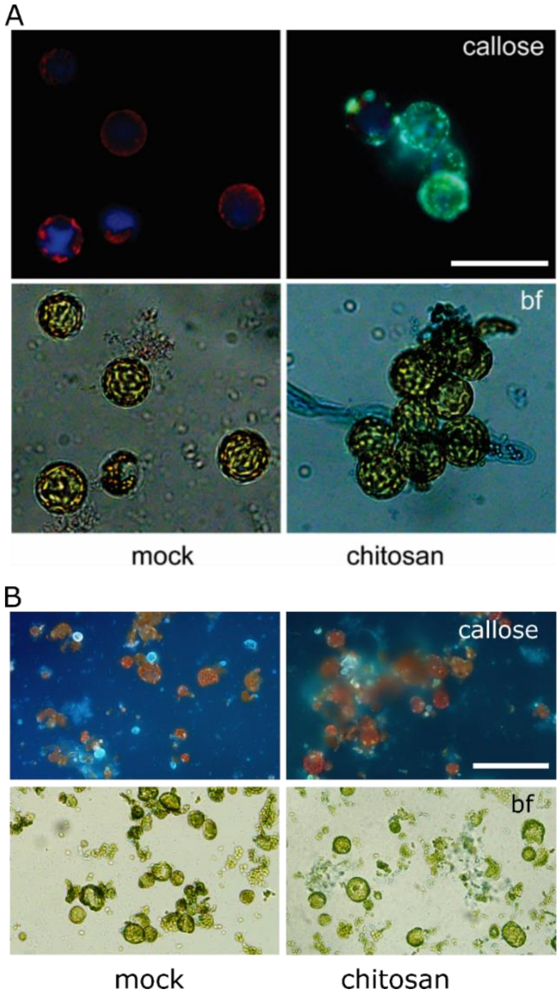


### Supplementary figure 5.

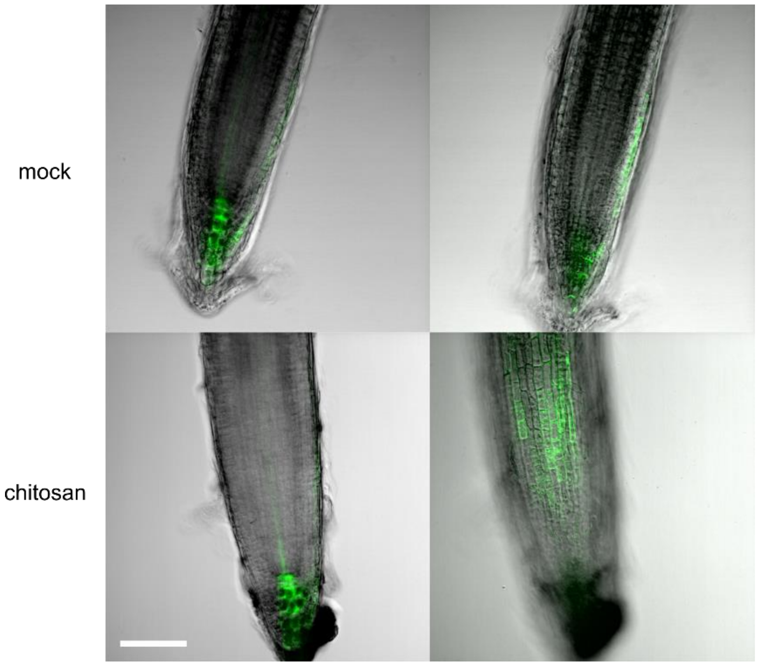
